# Ion-networks: a sparse data format capturing full data integrity of data independent acquisition mass spectrometry

**DOI:** 10.1101/726273

**Authors:** Sander Willems, Simon Daled, Bart Van Puyvelde, Laura De Clerck, Sofie Vande Casteele, Filip Van Nieuwerburgh, Dieter Deforce, Maarten Dhaenens

## Abstract

Data-independent acquisition (DIA) mass spectrometry (MS) has introduced deterministic, periodic and simultaneous acquisition of all fragment ions. Despite the chimeric side-effects associated with this unprecedented data integrity, DIA data analysis approaches still use conventional spectra and extracted ion chromatograms (XICs) that represent individual precursors and fragments. Here, we introduce ion-networks, an alternative data format wherein nodes correspond to reproducible fragment ions from multiple runs and edges correspond to consistent co-elution. Each ion-network represents a complete experiment and computationally eliminates chimericy based on reproducibility without sacrificing data integrity.

---

The last decade, liquid chromatography (LC)-MS techniques have been developed that remove the stochastic precursor selection of data-dependent acquisition (DDA). Thus, these DIA techniques acquire a reproducible periodic signal for each individual fragment of all eluting analytes. An early example of DIA is MS^e^, wherein each low energy (LE) precursor scan is followed by a single high energy (HE) fragment scan. Hereby a maximal data integrity is obtained for all precursors and their fragments [1]. However, acquisition with such high data integrity creates an additional challenge for data analysis in the form of chimericy: simultaneously acquired fragments rarely belong to the same precursor and cannot easily be related to each other or their original precursor anymore. This problem has been approached by perceiving DIA data as a combination of existing data formats. More specifically, there are examples of software tools that create DDA-like pseudo-spectra per precursor by retrieving their fragments [2]. Unfortunately, such static spectrum-centric representations inherently oversimplify DIA data. More commonly, targeted multiple-reaction-monitoring (MRM)-like approaches are taken that are based on XICs of individual ions, colloquially called peptide-centric in the proteomics domain [3, 4]. However, such targeted approaches are constrained by prior assumptions about sample content. In conclusion, loss of relevant fragment connectivity in a DIA run currently enforces analysis to revert to conventional LC-MS data formats.

Nowadays, DIA strategies reduce chimericy instrumentally to facilitate data analysis. The most widespread technique is cycling through the precursor mass-to-charge ratio (*m/z*) range with smaller yet more windows with e.g. sequential window acquisition of all theoretical mass spectra (SWATH) [5]. With the latest developments, these fixed sequential windows can even be replaced by a continuous quadrupole scanning such as scanningSWATH or scanning quadrupole DIA (SONAR) [6, 7]. Unfortunately, increasing selectivity requires either shorter scan times or an increased cycle time. The former results in reduced sensitivity while the latter gives poor periodic sampling and both reduce the duty cycle for any given ion, i.e. its percentage that reaches the detector. An orthogonal approach without any precursor selection and associated duty cycle loss, is to introduce additional separation with an ion mobility separation (IMS) cell before fragmentation as described in high definition MS^e^ (HDMS^e^) [8]. IMS, achieved in milliseconds, fits exactly between LC separation in seconds and time of flight (ToF) separation in microseconds. Hence, each ion can uniquely be defined by three coordinates: retention time (*t*_*R*_), drift time (*t*_*D*_) and *m/z*. However, accuracy of these coordinates is limited by resolution of instrumental separation and consequently not all chimericy can be resolved. More recently, IMS was combined with window selection in parallel accumulation serial fragmentation combined with data-independent acquisition (diaPASEF) [9]. Herein they exploit the relation between precursor *m/z* and *t*_*D*_ with quadrupole selection after IMS to maintain a high duty cycle for only those precursors of interest, at the cost of ignoring all others. In brief, regardless of the DIA technique employed there is an instrumental trade-off between data integrity and chimericy.

Here, we leverage DIA reproducibility to eliminate chimericy purely at the data level, redeeming data acquisition and data analysis from this effort. We collapse all runs from an entire experiment into a single noiseless ion-network prior to identification or quantification (Supplementary note 1, Figure SF1). The nodes of this ion-network are between-run aligned HE fragment ions and the edges represent consistent within-run co-elution. Herein noise is assessed by lack of reproducibility between runs. Equally, fragments from chimeric precursors are deconvoluted due to minor inconsistent stochastic differences between runs, while fragments from the same precursor exhibit consistent co-elution in each run as fragmentation occurs after precursor separation. Since the complete signal for each between-run reproducible HE fragment ion is collapsed into a single denoised and deconvoluted data point, the ion-network of a complete experiment becomes very sparse while retaining all relevant information from the acquired DIA data. This greatly simplifies fragment identification in e.g. untargeted analyses. Furthermore, the sparsity improves with the number of runs and is independent of the acquisition technique. In conclusion, acquisition and analysis are no longer directly connected and thus both can be developed unconstrained.

To illustrate the creation and characteristics of such an ion-network, we visualized an example with an interactive graphical browser (Supplementary note 1.8, Figure 1). This example is a public benchmark proteomic HDMSe dataset, which favors high data integrity at the cost of high chimericy. It contains a total of ten runs from two samples with different mixtures of tryptic Human, Yeast and E. coli peptides with organism mass fractions of respectively 1:1, 1:2 and 4:1, mimicking two different biological conditions [10] (Supplementary note 1.1.2). Peak picking at intensity threshold 1 and signal-to-noise ratio (SNR) 1 yielded on average ± 6,600,000 HE fragment ions per run (Supplementary note 1.2). After calibrating and aligning the *m/z, t*_*R*_ and *t*_*D*_ of all runs, ±3,000,000 (46%) HE fragment ions per run were found to be irreproducible noise, while ±540,000 (8%) were fully reproducible in all ten runs (Supplementary notes 1.4, 1.5.1, Figures 1A-B, SF2, SF3). As expected, the more reproducible an ion is, the higher its average intensity. These results indicate robust signal throughout four orders of magnitude and the capability to distinguish noise from signal. All (partially) reproducible ions can now be defined as nodes, i.e. aggregates, within our ion-network. For each aggregate pair, we set an edge if and only if they consistently co-elute within each run (Supplementary note 1.5.2, Figures 1C-D, SF4). Per aggregate in this particular ion-network, the median of consistently co-eluting aggregates is 24 with an interquartile range (IQR) of {6, 62}(Figure SF5). Of paramount importance; consistent co-elution is most evident between highly reproducible aggregates with similar intensity ratio profiles, i.e. derived from the same organism by benchmark design (Figures 1D, 1F, SF6). Thus, reproducibility can indeed deconvolute fragments from chimeric precursors.

**Figure 1:**
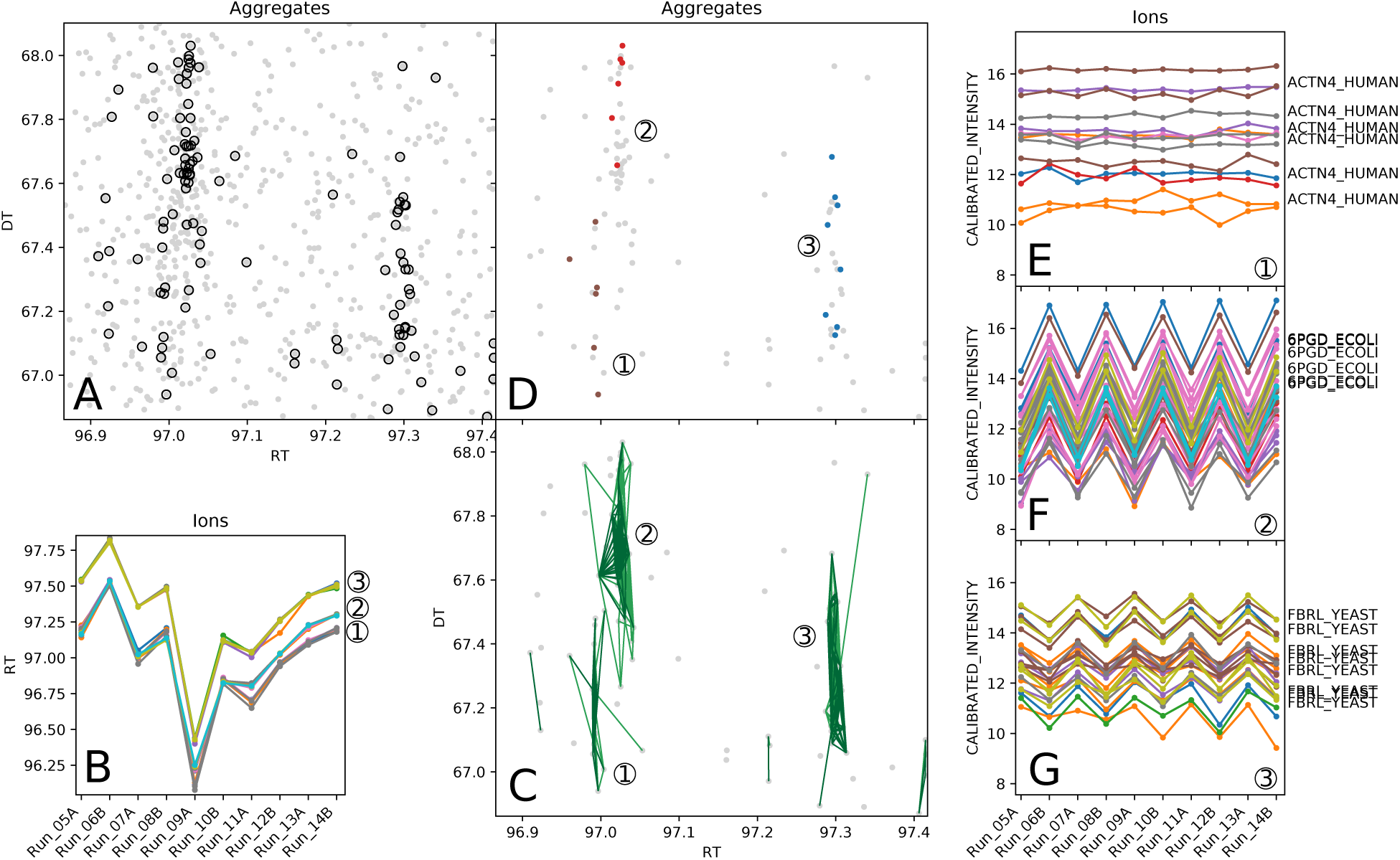
Ion-network example visualized with an interactive graphical browser. An ion-network was created and annotated for a total of ten public proteomic HDMS^e^ runs from two different benchmark samples. Herein the nodes **(dots in panels A, C-D)** are aggregates, i.e. between-run aligned HE fragment ions, while the edges **(green lines in panel C)** represent consistent within-run co-elution. With an interactive browser, we zoomed in on the aggregates of a select region, containing many aggregates with a reproducibility of at least 2 **(panel A)** including several that are fully reproducible **(circled dots in panel A, dots in panels C-D)**. After visualizing the *t*_*R*_ values of each ion per run for (a selection of) these fully reproducible aggregates **(panel B)**, three groups become apparent. Two of these groups co-elute in the first six runs, but are deconvoluted in the last four runs due to stochastic effects. For each potential pair of aggregates, an edge is set if and only if they consistently co-elute in each run, thereby forming the final network **(panel C)**. When this full network is annotated, multiple aggregates of three distinct peptides are annotated **(colored dots in panel D)**. Furthermore, when the individual ion intensities of the three clusters are visualized, each of their aggregates follow the same pattern indicating a correct deconvolution **(panels E-G)**. Notice that many unannotated aggregates, presumably other fragments that are not mono-isotopic singly charged b- or y-ions, still have a high quantification potential due to the deconvolution. Finally, the intensity patterns agree with the benchmark design wherein fragments from Human peptides **(1)**, E. coli peptides **(2)** and Yeast peptides **(3)** have an expected logarithmic fold change (logFC) of respectively 0, -2 and 1, thereby confirming correct identification.

To demonstrate the applicability of ion-networks on the analysis side, we implemented a simplistic untargeted database search algorithm to annotate individual aggregates, i.e. reproducible HE fragment ions, in the proteomics domain (Supplementary note 1.7). Conceptually, this results in a peptide-fragment-to-ion-neighborhood match (PIM) for an aggregate in a similar way as a precursor in DDA has a peptide-to-spectrum match (PSM). To illustrate the performance on our benchmark ion-network, we annotated the aggregates with singly-charged mono-isotopic b- and y-fragments from tryptic Human, Yeast and E. coli peptides without missed cleavages or variable modifications. Hereby ±99,000 aggregates were annotated, belonging to ±8,900 unique peptide sequences of ±2,100 unique protein groups, all at their respective 1% false discovery rate (FDR) (Figures 1E-G). Notably, the intensity ratios of the annotated aggregates coincide with expected organisms demonstrating a correct FDR estimation (Figure SF7). This simplistic untargeted database search algorithm greatly enriched fully reproducible aggregates, again indicating the power of DIA reproducibility to denoise and deconvolute (Figure SF8).

On the acquisition side, we demonstrate the versatility and performance of ion-networks with several experimental designs and acquisition strategies (Supplementary note 1.3). To this end, we first created an annotated ion-network of a publicly available SONAR HeLa dataset of nine runs from a single sample. Herein we imported the scanning quadrupole selection as if it was *t*_*D*_, showing the versatility of ion-networks to modify this dimension according to the data at hand. Without any other adjustments, this resulted in significant annotations at 1% FDR for ±29,000 aggregates, ±5,800 unique peptide sequences and ±1,600 protein groups. Second, we created a mock ion-network for a single DDA ToF run from a sample containing three mixed proteomes. This mock ion-network is a list of all acquired HE fragment ions grouped in fully connected clusters per tandem-MS spectrum. Herein denoising was done upfront through peak picking with default intensity-based SNR filtering instead of between-run reproducibility. Deconvolution was done upfront by quadrupole selection instead of consistent within-sample co-elution. With our simplistic annotation approach, we were able to annotate ±160,000 ions, ±5,900 peptides and ±1,500 proteins at 1% FDR. Surprisingly, 22% of the edges within this network are between ions with different peptide annotations, which is higher than all DIA ion-networks that only have 3-9% chimeric edges. From this perspective, reproducibility and consistent co-elution can outperform DDA quadrupole selection in terms of reducing noise and chimericy (Figure SF9 and Table ST1). Together, these results imply that ion-networks are most performant on datasets with high data integrity as well as high chimericy. To confirm this hypothesis, we acquired precursorless HDMS^e^ which we denominated single window ion mobility (SWIM)-DIA. With only a single continuously acquired HE scan and no precursor window selection, SWIM-DIA has the highest possible data integrity for fragment ions. To illustrate its performance, we created three ion-networks for an in-house benchmark dataset with a similar design as Navarro et al. [11]. These ion-networks comprise a set of HDMS^e^ runs, a set of SWIM-DIA runs and a set with runs from both acquisitions combined, all from the same samples (Supplementary note 1.1.1). The median coefficient of variation (CV) of fully reproducible aggregates reduces from 15.1% in HDMS^e^ to 12.7% in SWIM-DIA without any annotation, let alone summarization, illustrating an unprecedented quantitative accuracy (Figure SF10).

In conclusion, ion-networks are able to capture HE fragment ions from different DIA techniques in a very sparse format with minimal noise and chimericy. While we only investigated a single software application, i.e. a simplistic proteomic database search, we postulate that the noiseless nature of these ion-networks enables a plethora of other untargeted software applications such as e.g. proteomic *de novo* algorithms, metabolomics database searches, quantification-centered workflows, et cetera. Additionally, ion-networks enabled a novel hardware application with maximal data integrity and an unprecedented quantitative accuracy, i.e. SWIM-DIA.

## Acknowledgements

This research was primarily funded by the Research Foundation Flanders (FWO) through research project grant G013916N, mandate 12E9716N (MD) and mandate 11B4518N (BVP), as well as Flanders Innovation & Entrepreneurship (VLAIO) mandate SB-141209 (LDC). We thank Hans Vissers, Scott Geromanos, Steve Cievarini (Waters Corporation) and Lennart Martens (VIB, Ghent, Belgium) for their critical feedback. LC-MS runs were acquired at the ProGenTomics facility and computational assistance was provided by Yannick Gansemans and Laurentijn Tilleman (Ghent University, Ghent, Belgium).

## Author contributions

SW and MD conceived the idea of creating noiseless ion-networks with reproducibility for untargeted DIA analysis. SW, SD and MD envisioned SWIM-DIA as hardware application. SW performed all computational analysis. SD and BVP performed all sample preparation and data acquisition. MD and DD supervised the project. SW and MD wrote the draft manuscript. All authors provided critical feedback during research and writing.

## Supplementary figures and tables

**Figure SF1:**
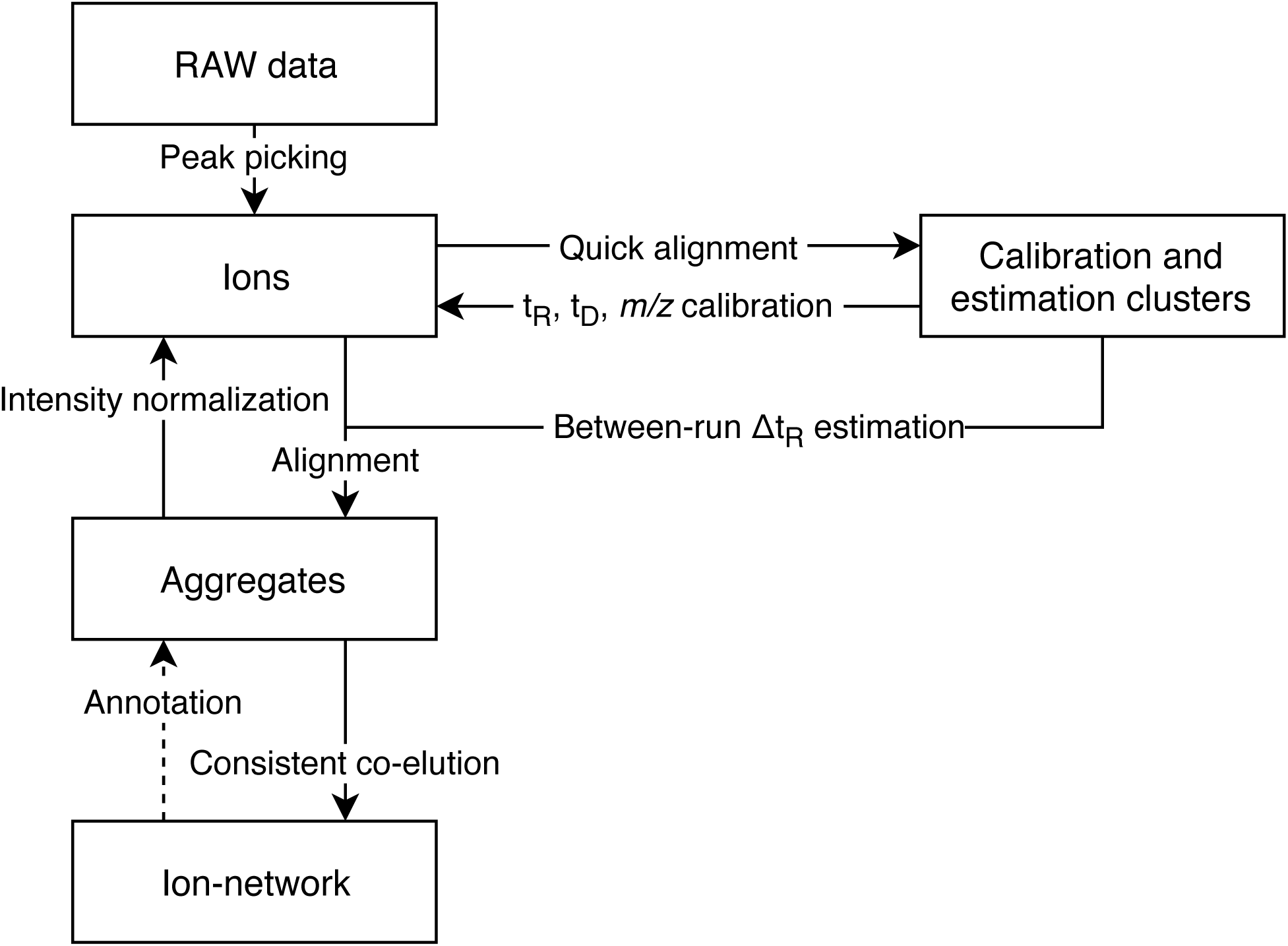
Schematic overview of the creation of an ion-network. Raw liquid chromatography (LC)-mass spectrometry (MS) data. (Supplementary note 1.1) from an experiment with multiple runs from different samples (Supplementary note 1.3) are the start of creating a noiseless ion-network. First, all runs are **peak picked** (Supplementary note 1.2) to obtain an exhaustive list of **ions** that can be analyzed concurrently. Herein each peak picked ion has a mass-to-charge ratio (*m/z*) apex, drift time (*t*_*D*_) apex, retention time (*t*_*R*_) apex and run identifier as primary coordinates, as well as meta-data describing the intensity and apex peak picking errors per coordinate. The 50,000 most abundant ions per run are used in a **quick alignment** to determine which are fully reproducible in all runs. These fully reproducible ions form **clusters** that are used to **calibrate** the primary coordinates between each run and furthermore give an **estimate** of the between-run deviation of the *t*_*R*_ (Supplementary note 1.4). Based on these calibrated coordinates, all ions from the complete experiment are **aligned** into **aggregates**, i.e. between-run reproducible ions (Supplementary note 1.5.1). With these aggregates, the intensity of each ion is **normalized** per run (Supplementary note 1.6). Next, aggregates with at least two constituent ions are defined as nodes in the **ion-network**, while irreproducible ions are considered noise and discarded. For each pair of aggregates, an edge is set if and only if their constituent ions **consistently co-elute** within each run (Supplementary note 1.5.2). Hereby fragments from chimeric precursors can be deconvoluted, as stochastic co-elution of precursors is not always consistent. For proteomics experiments, each individual aggregate within this ion-network can be **annotated** as a specific b- or y-ion with a simplistic database search (Supplementary note 1.7).

**Figure SF2:**
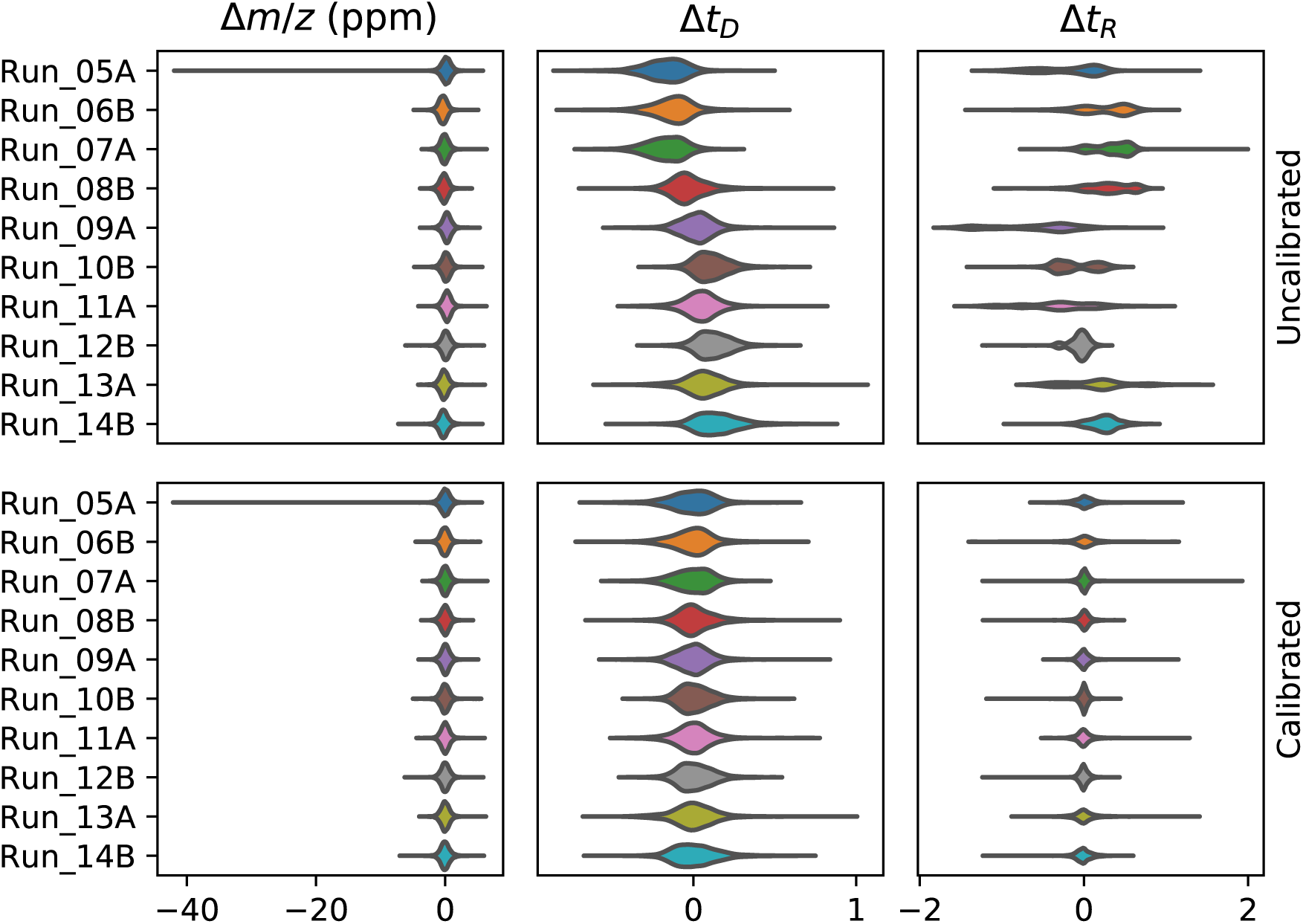
Between-run calibration. For all high definition MS^e^ (HDMS^e^) runs of from PXD001240 (*y***-axis)**, the 50,000 most abundant ions after peak picking are selected for a quick alignment. Herein, clusters of exactly ten ions, each from a different runs, are detected in the *m/z* space and clusters with outliers in the *t*_*R*_ and *t*_*D*_ space are removed. The remaining 5,270 clusters containing aligned ions of each run are equally partitioned over a calibration- and validation set. For each calibration cluster, the distance of its aligned ions in *m/z* (in parts per million (ppm)), *t*_*D*_ and *t*_*R*_ to the cluster average is determined per run **(top)**. Hereafter, the *m/z, t*_*D*_ and *t*_*R*_ of all ions are calibrated per run (Supplementary note 1.4). To estimate the performance of this calibration, the distance in calibrated *m/z* (in ppm), *t*_*D*_ and *t*_*R*_ between the validation cluster averages and their aligned ions is determined per run **(bottom)**.

**Figure SF3:**
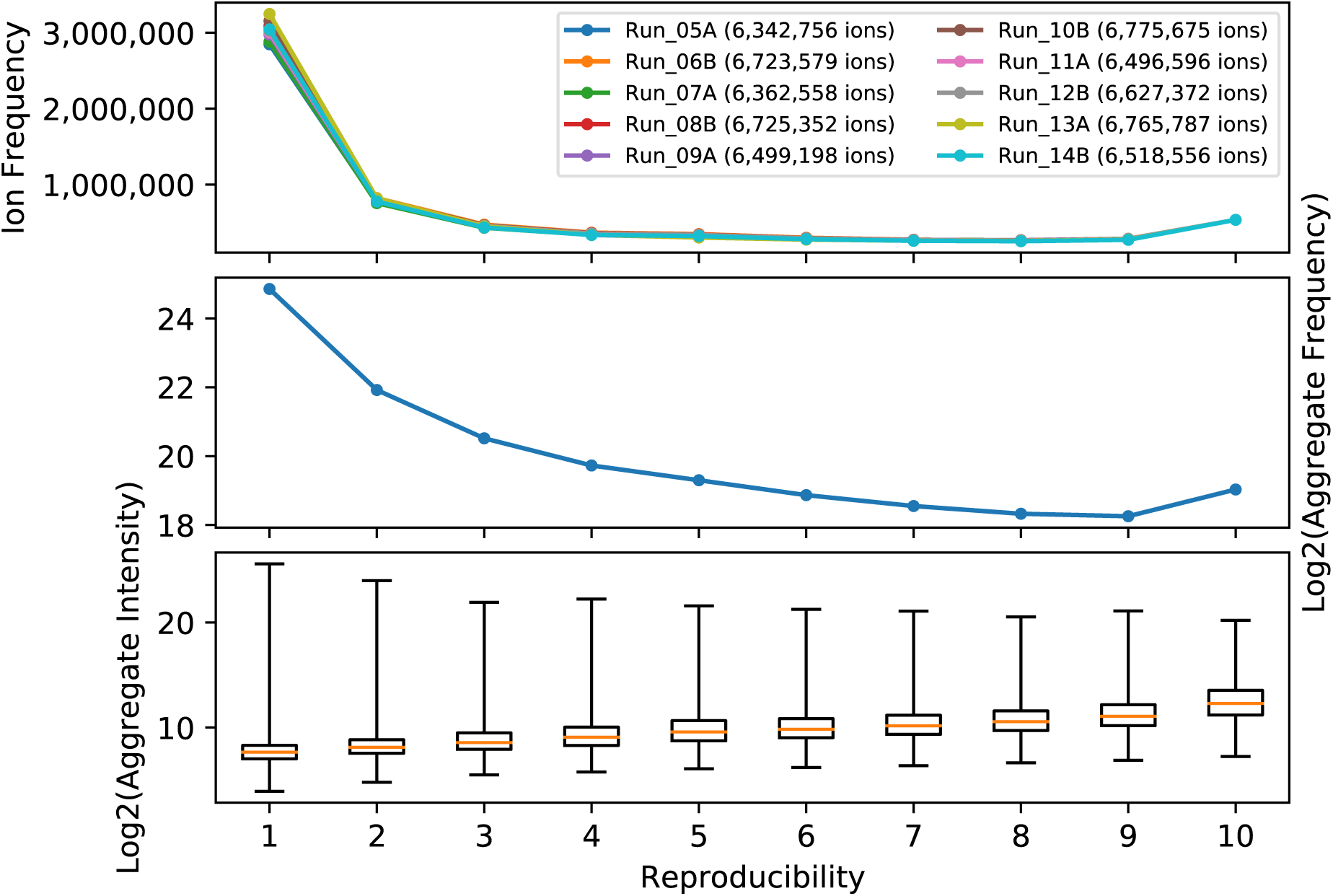
Aggregate counts and intensities. After peak picking and calibration, all 65,837,429 ions from all HDMS^e^ runs from PXD001240 **(top legend)** are aligned into aggregates (Supplementary note 1.5.1). An aggregate is defined as a set of unique ions from different runs with equal calibrated *m/z, t*_*D*_ and *t*_*R*_, wherein the number of different runs is expressed as the reproducibility of an aggregate. As such, each ion of a specific run is contained in exactly one aggregate **(top)**. Irreproducible aggregates with only one ion are retained here only to illustrate the amount of noise, but are discarded in subsequent analyses. For each of the 39,361,063 aggregates **(middle)**, the average intensity of its ions was calculated, irrespective of whether the ions came from sample A or B **(bottom)**. Boxplots indicate interquartile range (IQR) with median **(orange line)** while whiskers extend to the minima and maxima.

**Figure SF4:**
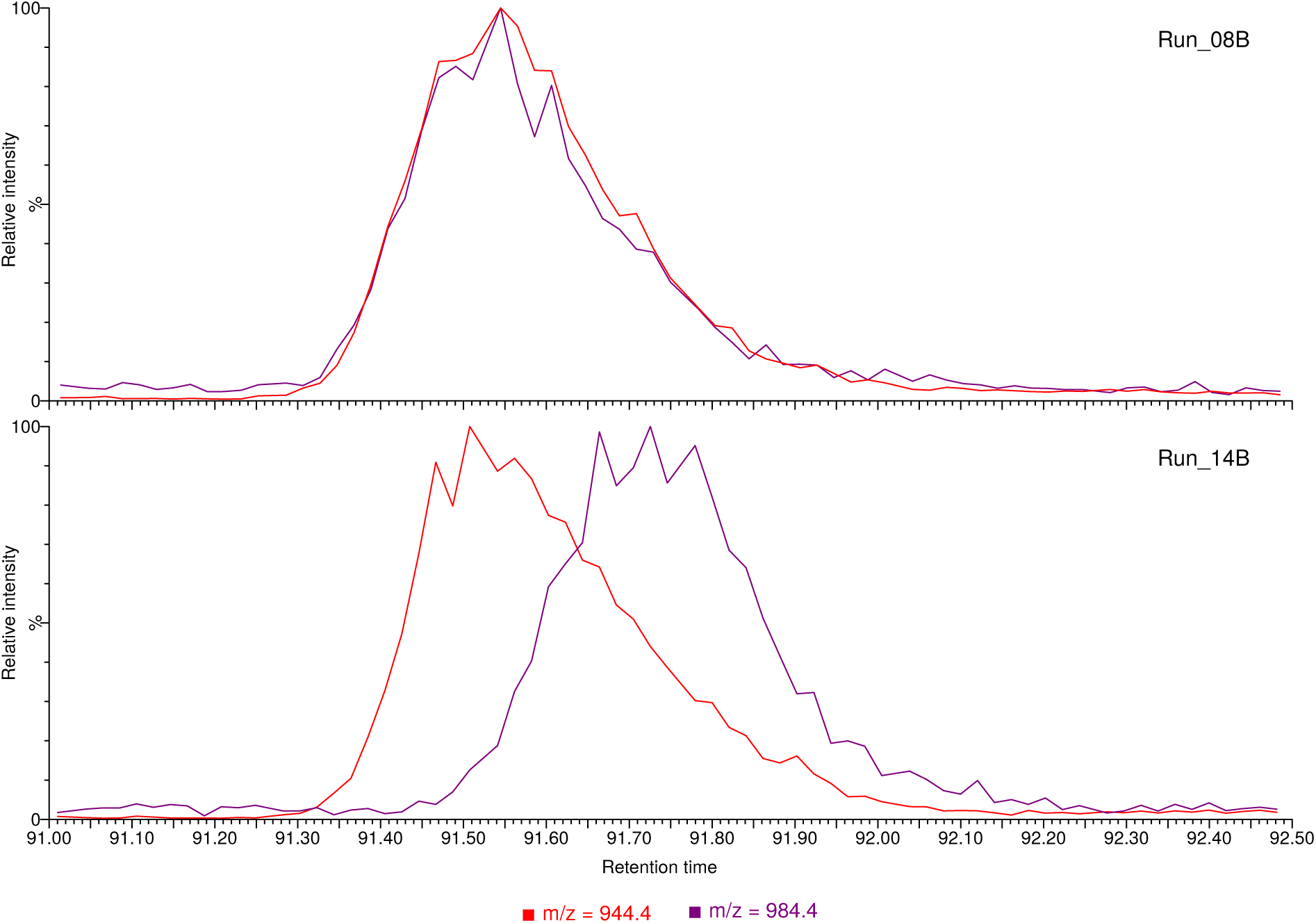
Example of non-consistent co-elution. Two high energy (HE) fragment ions with *m/z* 944.4 **(red)** and 984.4 **(purple)** are co-eluting in run 8 **(top)** of PXD001240, with equal *t*_*R*_ apices and similar peak shapes. However, in run 14 **(bottom)** of the same dataset, their *t*_*R*_ apices are separated by several seconds and different peak shapes, making it unlikely that these HE fragment ions originate from the same low energy (LE) precursor ion. This hypothesis is confirmed by their intensity ratio profiles revealing these HE fragment ions belong to different organisms by design of the benchmark. When both runs are analyzed simultaneously with an ion-network, this inconsistent co-elution can be leveraged to deconvolute the two chimeric HE fragment ions from run 8.

**Figure SF5:**
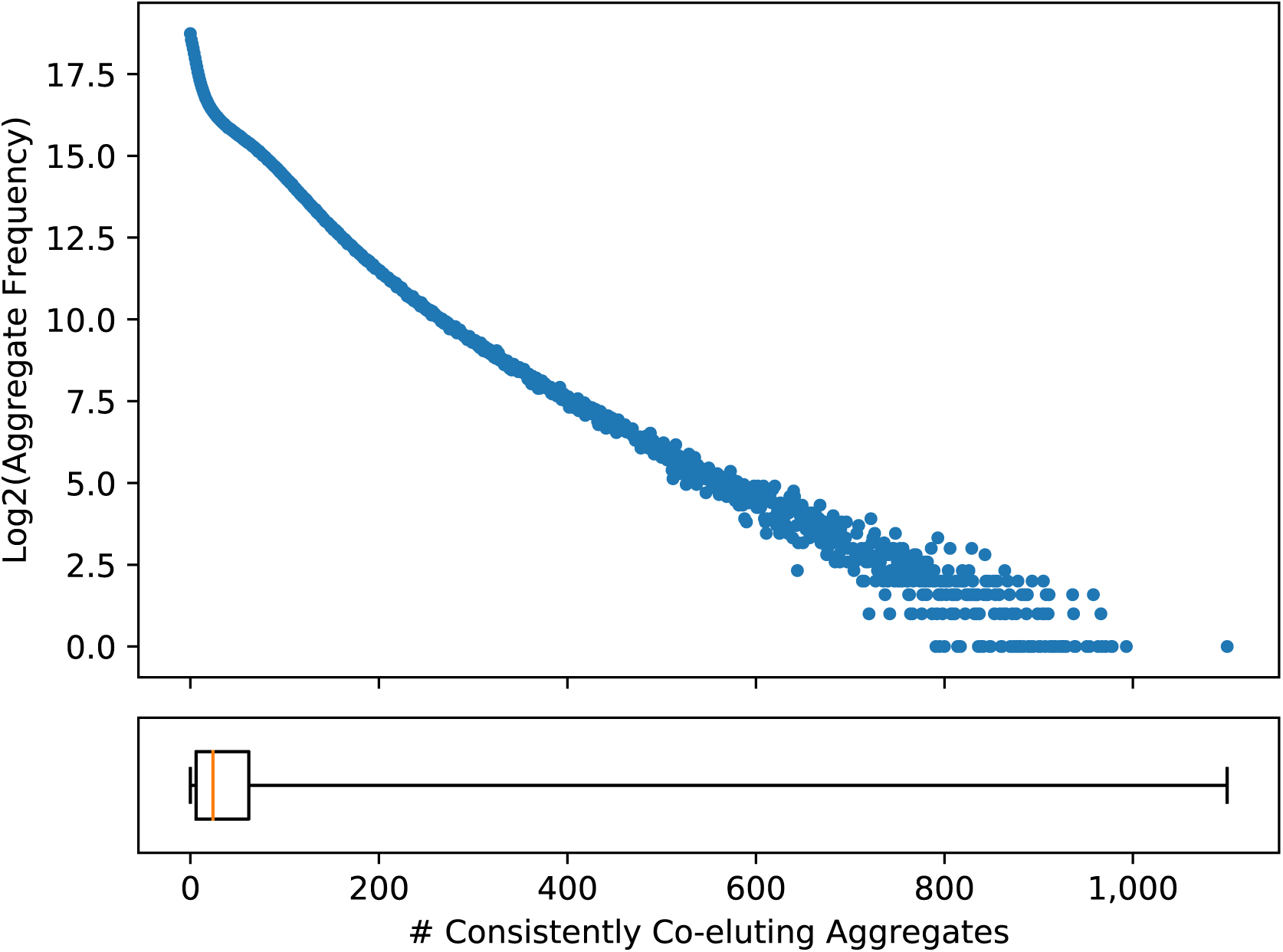
Consistently co-eluting aggregates. An HDMS^e^ ion-network was created for PXD001240 (Supplementary note 1.5). Herein, each aggregate, i.e. (partially) reproducible HE fragment ion, has a number of consistently co-eluting aggregates (*x***-axis)** and the logarithmic frequencies of these aggregates with equal consistently co-eluting aggregates (*y***-axis)** was determined. Consistently co-eluting aggregates are presumed to originate from the same LE ion and comprise all potential HE fragment ions such as b- and y-ions, isotopes, neutral losses, et cetera. The boxplot indicates the IQR with a median **(orange line)** of 24 and whiskers extending to the minima and maxima.

**Figure SF6:**
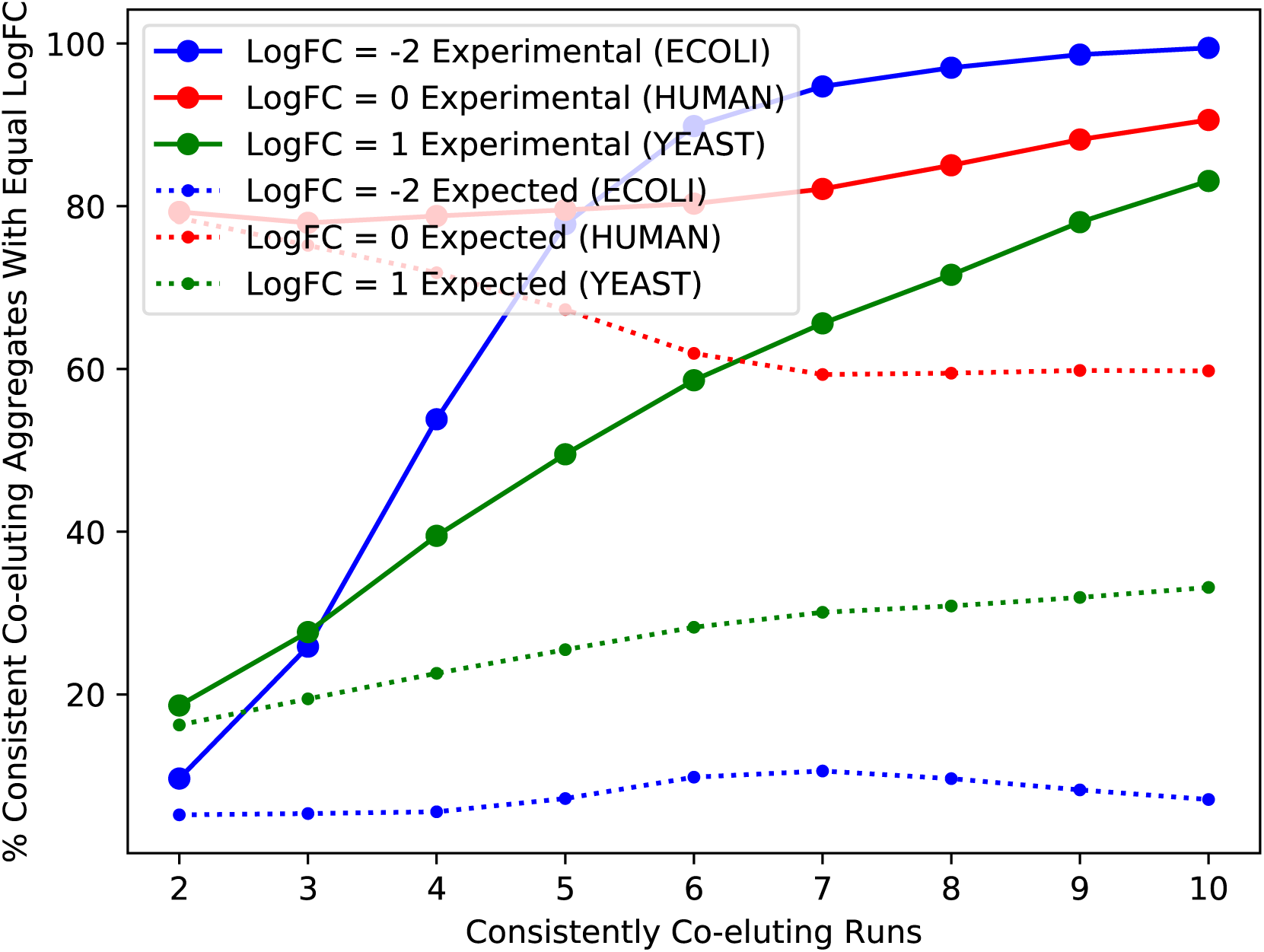
Intensity ratios of consistently co-eluting aggregates. An ion-network was created for public dataset PXD001240 (Supplementary note 1.5). This benchmark contains five HDMS^e^ runs from a sample in condition A and five runs from a sample in condition B, each consisting of a mixture of Human, Yeast and E. coli tryptic peptides with mass fractions of respectively 65/15/20 and 65/30/5. If an aggregate in this ion-network contains ions from runs of both condition A and B, the logarithmic fold change (logFC) of this aggregate can be determined. By design of the benchmark, aggregates with a logFC of 0, 1 or -2 are respectively expected to be Human **(red)**, Yeast **(green)** or E. coli **(blue)**. When all pairs of aggregates are partitioned by the number of runs in which they consistently co-elute (*x***-axis)**, the percentage of paired aggregates with equal logFC (*y***-axis)**, i.e. likely organism origin, can be determined **(experimental; full lines)**. While an equal organism origin does not proof that the pair of aggregates are fragments from the same precursor, the converse statement is generally true: a pair of aggregates with different logFC are fragments from two different chimeric precursors that are not deconvoluted. To determine the impact of consistent co-elution on this deconvolution, we calculated the theoretical probability that a pair of aggregates has the same logFC **(expected; dotted lines)**, regardless of consistent co-elution. This was done by first calculating the probability *P* (*X*) of an aggregate for *X* ∈ {Human, Yeast, E. coli} per partition of consistent co-elution. When aggregates within these partitions are paired independently, a pair has the same logFC with probability *P* (both *X*) = *P* (*X*)^2^.

**Figure SF7:**
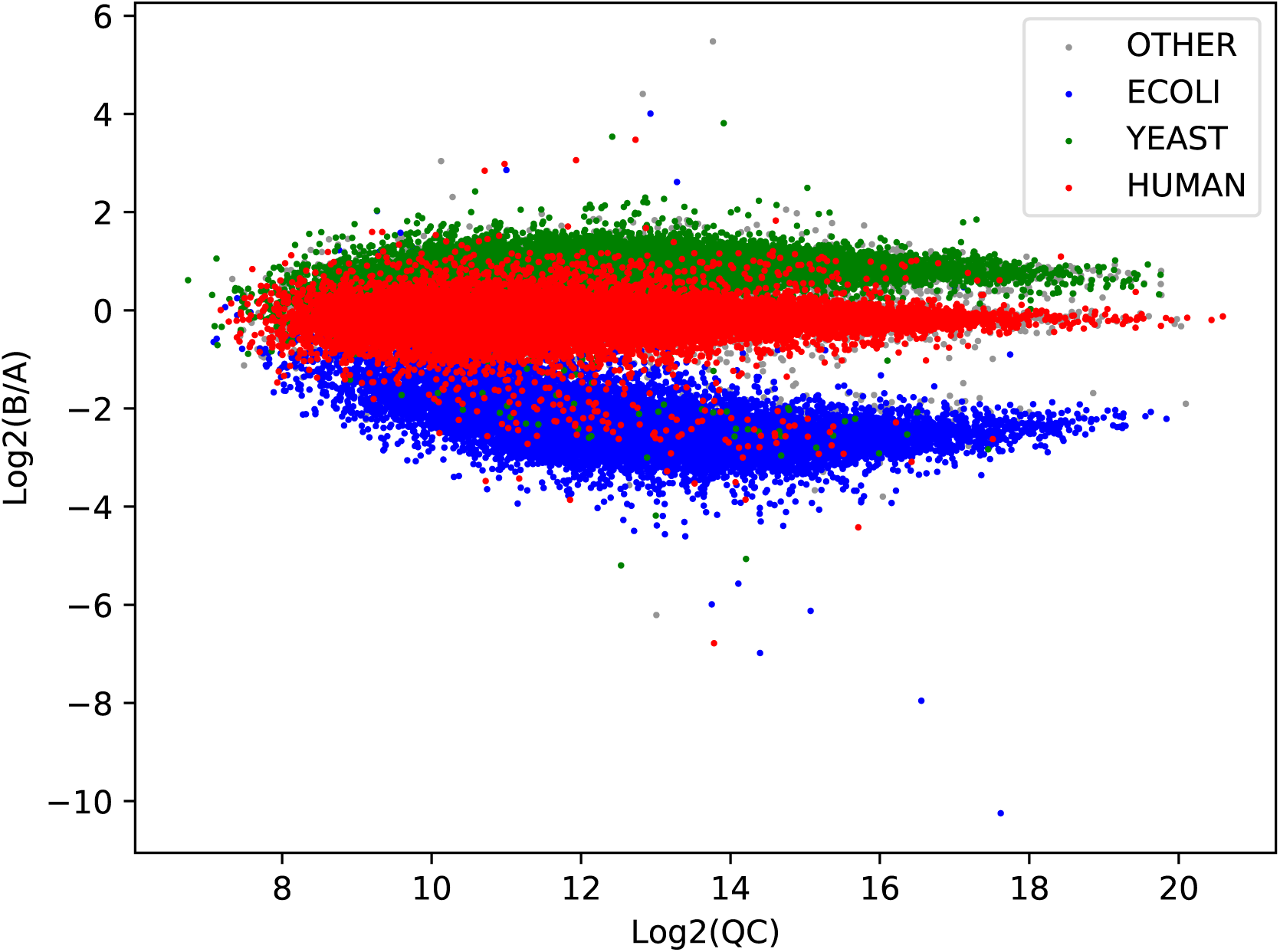
logFCs of annotated aggregates. 99,375 aggregates, i.e. (partially) reproducible HE fragment ions, in the HDMS^e^ ion-network of public dataset PXD001240 were annotated following a simplistic untargeted database search (Supplementary note 1.7). Within this benchmark dataset, Human **(red)**, Yeast **(green)** and E. coli **(blue)** have expected logFCs (*y***-axis)** of respectively 0, 1 and -2. Other annotations **(grey**) include peptide sequences from the common repository of adventitious proteins (cRAP) and peptide sequences assignable to multiple proteins. To estimate the accuracy of each logFC calculation, the average intensity (*x***-axis)** of each aggregate was determined by taking the unweighted average of condition A and B.

**Figure SF8:**
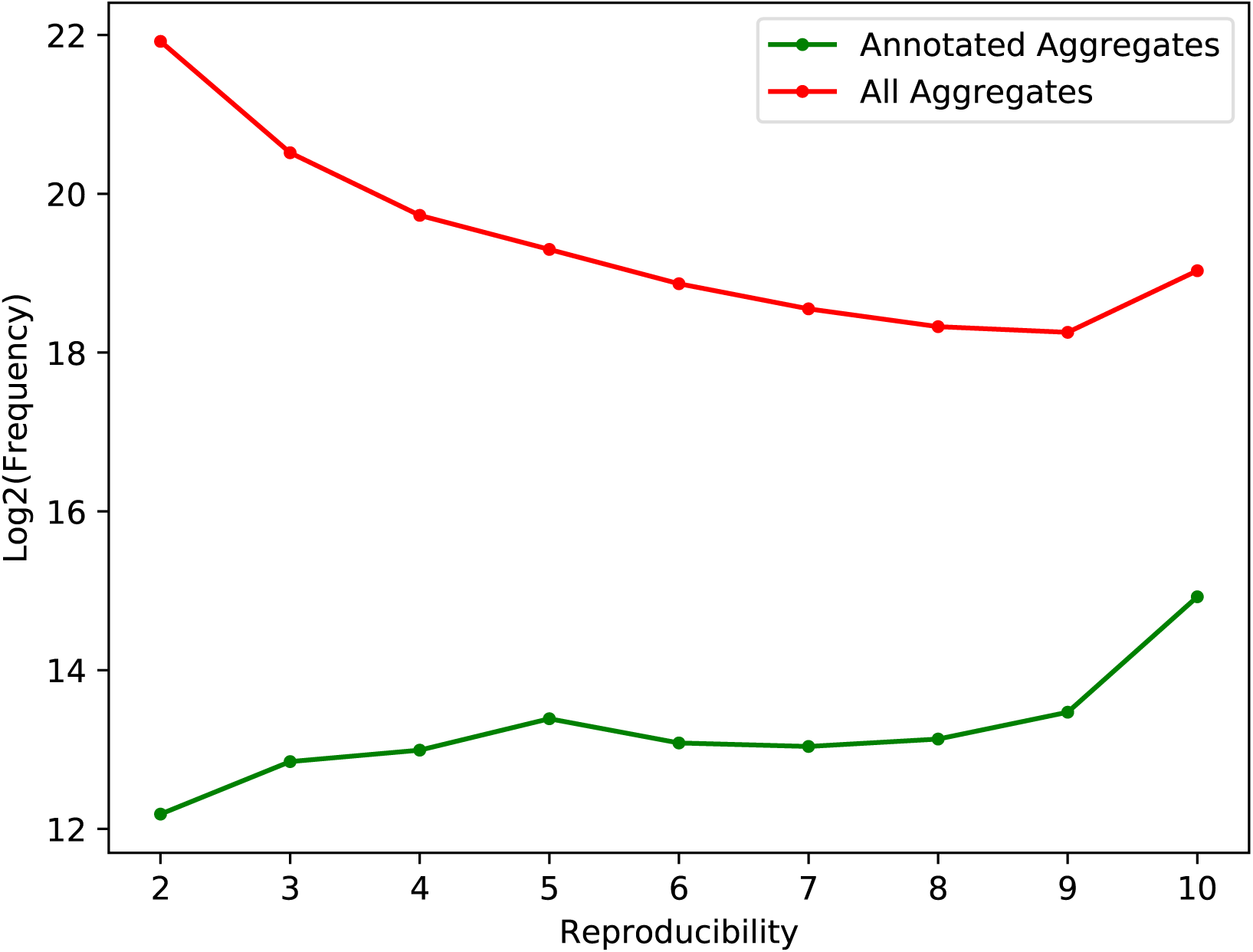
Annotated aggregate frequencies. Within the HDMS^e^ ion-network of PXD001240, 99,375 aggregates were annotated with a significant score **(green)** (Supplementary note 1.7). The logarithmic frequency (*y***-axis)** of these aggregates was determined in function of their reproducibility (*x***-axis)**. This was compared against the logarithmic frequency and reproducibility of all aggregates in the whole ion-network **(red)**, regardless of their annotation. Hereby annotation efficiency seems to be related to reproducibility as e.g. only 0.1% of two-fold reproducible aggregates were annotated, while 6% of all fully reproducible aggregates were annotated.

**Figure SF9:**
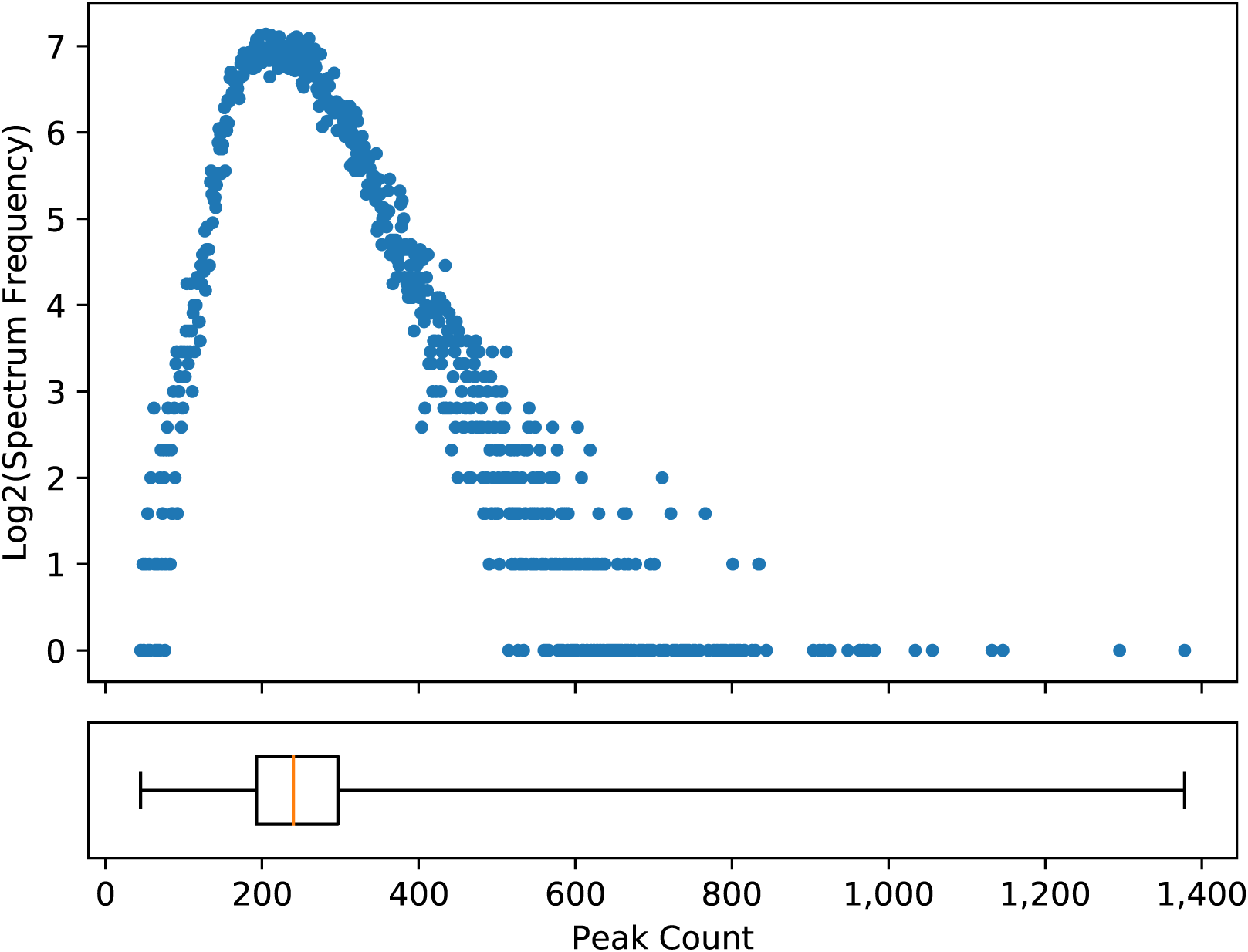
Number of peaks in a data-dependent acquisition (DDA) time of flight (ToF) spectrum. For a quality control (QC) sample containing a mixture of 65% Human, 22.5% Yeast and 12.5% E. coli, a single DDA ToF run was acquired. After peak picking with default intensity-based signal-to-noise ratio (SNR) filtering using Progenesis QI for Proteomics (Nonlinear Dynamics), 23,059 HE spectra were obtained. The logarithmic frequency (*y***-axis)** of the number of peaks (*x***-axis)** in these spectra was determined. The boxplot indicates the IQR with a median **(orange line)** of 240 and whiskers extending to the minima and maxima.

**Table ST1:**
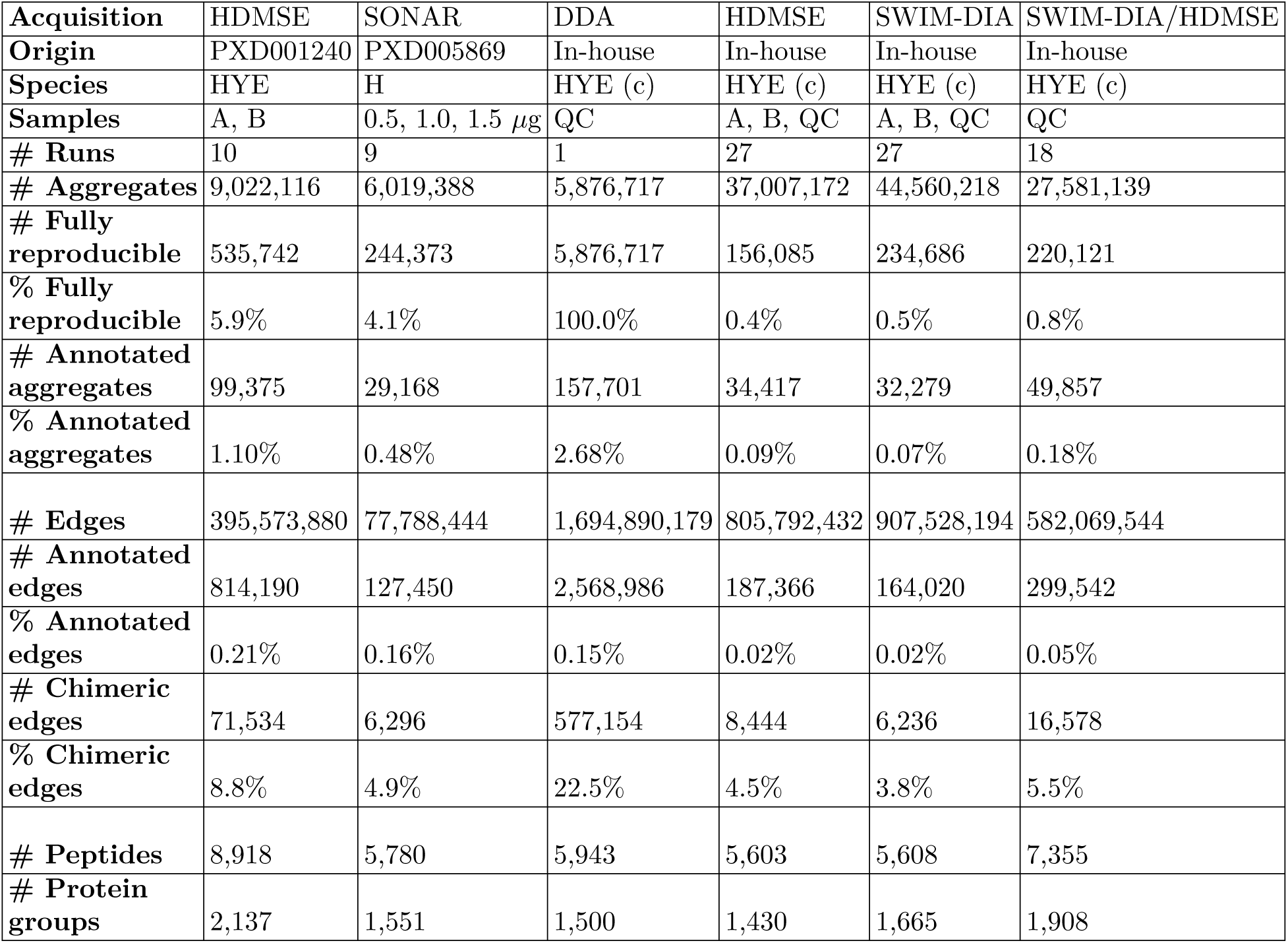
Statistics of ion-networks from different acquisitions. Six different ion-networks were analyzed. These ion-networks comprise very different datasets in terms of acquisition, species **((H)uman, (Y)east and (E). coli, from both (c)ommerical and non-commercial digests)** and samples **(mass fractions of A: 65/15/20, B: 65/30/5 and QC: 65/22.5/12.5)**, and number of runs (Supplementary note 1.3). Even so, all of these ion-networks were created and annotated with identical parameters. Annotated edges are defined as an edge between two annotated aggregates, while chimeric edges are defined as an edge between two aggregates with different annotated peptide sequences, where leucine and isoleucine were considered identical.

| Acquisition | HDMSE | SONAR | DDA | HDMSE | SWIM-DIA | SWIM-DIA/HDMSE |
| --- | --- | --- | --- | --- | --- | --- |
| Origin | PXD001240 | PXD005869 | In-house | In-house | In-house | In-house |
| Species | HYE | H | HYE (c) | HYE (c) | HYE (c) | HYE (c) |
| Samples | A, B | 0.5, 1.0, 1.5 μg | QC | A, B, QC | A, B, QC | QC |
| # Runs | 10 | 9 | 1 | 27 | 27 | 18 |
| # Aggregates | 9,022,116 | 6,019,388 | 5,876,717 | 37,007,172 | 44,560,218 | 27,581,139 |
| # Fully reproducible | 535,742 | 244,373 | 5,876,717 | 156,085 | 234,686 | 220,121 |
| % Fully reproducible | 5.9% | 4.1% | 100.0% | 0.4% | 0.5% | 0.8% |
| # Annotated aggregates | 99,375 | 29,168 | 157,701 | 34,417 | 32,279 | 49,857 |
| % Annotated aggregates | 1.10% | 0.48% | 2.68% | 0.09% | 0.07% | 0.18% |
| # Edges | 395,573,880 | 77,788,444 | 1,694,890,179 | 805,792,432 | 907,528,194 | 582,069,544 |
| # Annotated edges | 814,190 | 127,450 | 2,568,986 | 187,366 | 164,020 | 299,542 |
| % Annotated edges | 0.21% | 0.16% | 0.15% | 0.02% | 0.02% | 0.05% |
| # Chimeric edges | 71,534 | 6,296 | 577,154 | 8,444 | 6,236 | 16,578 |
| % Chimeric edges | 8.8% | 4.9% | 22.5% | 4.5% | 3.8% | 5.5% |
| # Peptides | 8,918 | 5,780 | 5,943 | 5,603 | 5,608 | 7,355 |
| # Protein groups | 2,137 | 1,551 | 1,500 | 1,430 | 1,665 | 1,908 |

**Figure SF10:**
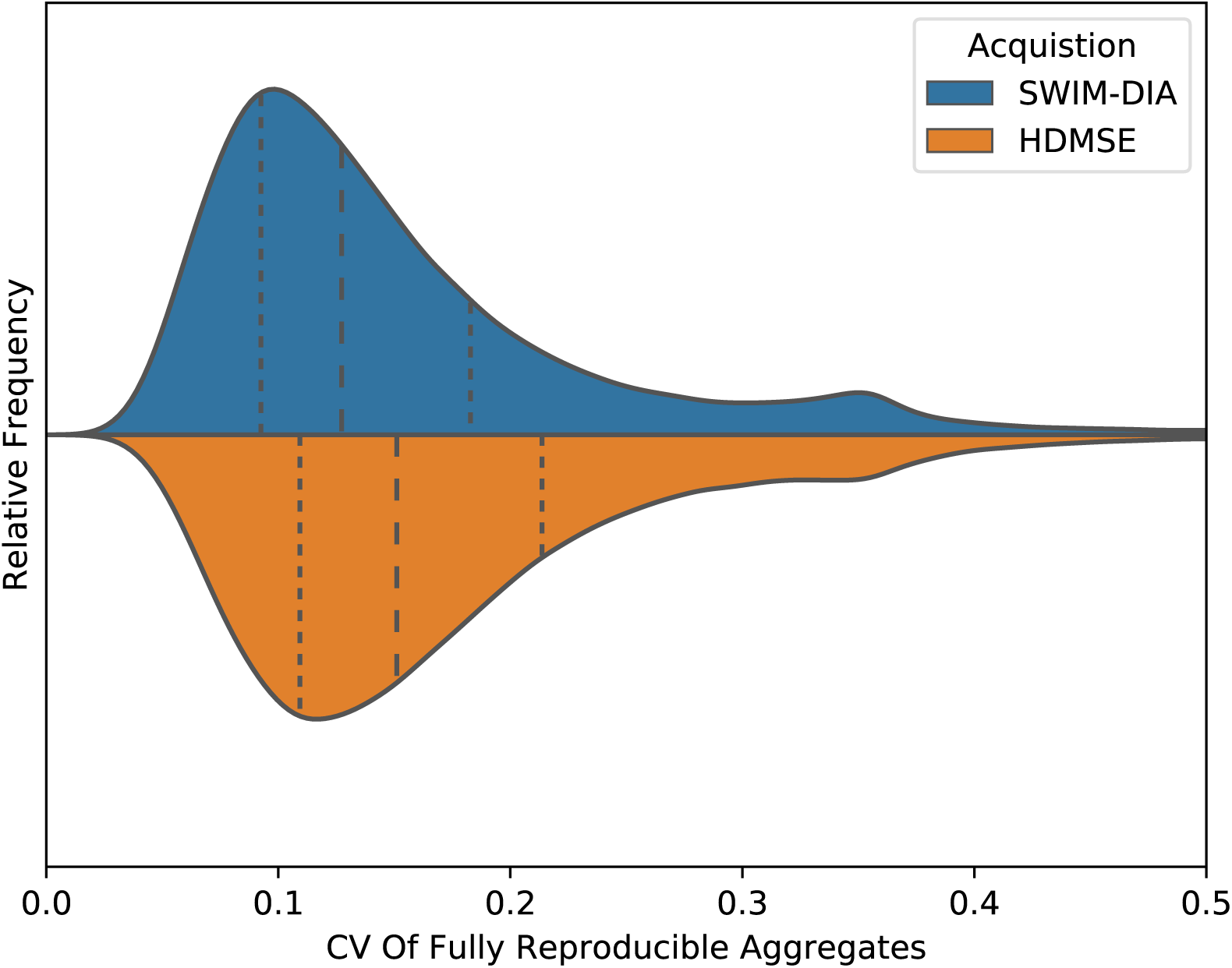
Distribution of the coefficient of variation (CV) values of single window ion mobility (SWIM)-data-independent acquisition (DIA) and HDMS^e^ aggregate quantification. For a QC sample of a mixture of commercial Human, Yeast and E. coli tryptic peptides (Supplementary note 1.1), nine runs were acquired in turn for both SWIM-DIA and HDMS^e^ mode. They were analyzed simultaneously to create a single ion-network with normalized intensities (Supplementary notes 1.5, 1.6). Hereby 220,121 fully reproducible aggregates were found. The CV (*x***-axis)** of these aggregates in the nine SWIM-DIA replicates **(top)** and the nine HDMS^e^ replicates **(bottom)** was determined, as well as their first, second and third quantiles **(dashed lines)**. Manual inspection of aggregates with CV around 0.35 for both SWIM-DIA and HDMS^e^ frequently shows aggregates with an outlying ion in the *m/z, t*_*R*_ or *t*_*D*_ dimension, which presumably is poorly aligned. A paired *t*-test shows there is a significant (*p* ≪ 10^−300^) difference between the CV of SWIM-DIA and HDMS^e^.

## Supplementary note

### 1 Material and methods

A brief overview of how all chapters in this material and methods section connect to each other is given in Supplementary Figure SF1.

#### 1.1 Raw data

Within this manuscript, raw mass spectrometry (MS) data from multiple samples were used.

##### 1.1.1 In-house samples

###### Sample preparation

Human, Yeast and E. coli commercial digests (carbamidomethylated tryptic peptides) were obtained from Promega (V6951, V7461) and Waters Corporation (186003196) respectively. The lyophilized samples were resuspended in 0.1% formic acid and spiked with 11 standard iRT peptides (Biognosys) in a 1/20 ratio. Two mixtures were prepared with different mass fractions; mixture A with 65% Human, 15% Yeast, and 20% E. coli and mixture B with 65% Human, 30% Yeast, and 5% E. coli. Thus, the resulting mixtures contain the same peptides with a logarithmic fold change (logFC) of 0, 1 and -2 for the respective organisms. Both mixtures were combined in equal volumes to create a quality control (QC) sample with 65% Human, 22.5% Yeast and 12.5% E. coli.

###### Data acquisition

Nine runs of 500 ng for each sample (A, B and QC) were acquired on a Synapt G2-Si (Waters Corporation) in both high definition MS^e^ (HDMS^e^) and single window ion mobility (SWIM)-data-independent acquisition (DIA). A default E. coli AutoQC sample was analyzed per nine runs to assess instrumental MS performance. HDMSe alternates between a low energy (LE) precursor scan and a high energy (HE) fragment scan, while SWIM-DIA only acquires HE fragment scans. Accumulation times and mass range for all scans were set to 600 ms and 50-2000 mass-to-charge ratio (*m/z*), which results in a cycle time of 1200 ms for HDMS^e^ and 600 ms for SWIM-DIA, ignoring between-scan times. For the HE fragment scans, collision energy was ramped from 10 to 60 V through the ion mobility separation (IMS) to apply optimal collision energies to the precursor ions [1]. The Synapt G2-Si was interfaced to a nano-acquity liquid chromatography (LC) system (Waters Corporation), operated under nano-conditions without trapping. Runs were separated on a C18 M-class column, 7 *µ*m x 250 mm with a 1.8 *µ*m particle size (Waters Corporation, 186007474). Flowrate was set to 300 nL/min with a 150 minute gradient (2-40 %B). The composition of mobile phase A was 0.1% formic acid with 3% DMSO in water and B was 0.1% formic acid in acetonitrile (percentages expressed in mass fractions). Finally, a single run of a QC sample was acquired in data-dependent acquisition (DDA) mode with equal LC settings.

##### 1.1.2 Public samples

Raw data were downloaded from ProteomeXchange for all ten HDMS^e^ runs of PXD001240 [2]. In brief, these data were assembled similar to the in-house samples to generate five runs for two conditions (A and B) with logFCs of 0, 1 and -2 for respectively Human, Yeast and E. coli. Note however that Human and Yeast peptides were obtained through non-commercial digests from alternative cell-lines. As such, these public samples are incomparable with our in-house samples of commercial digests.

Raw data were downloaded from ProteomeXchange for all nine scanning quadrupole DIA (SONAR) runs of PXD005869 [3]. In brief, a HeLa Human cell line was digested and three runs for three different sample amounts (0.5 *µ*g, 1.0 *µ*g and 1.5 *µ*g) were acquired for this single sample.

#### 1.2 Peak picking

Raw data from both public and in-house runs were peak picked with Apex3D (Waters Corporation) version 3.1.0.9.5. Parameters were set to a lockMass of 785.8426 for charge 2 with *m/z* tolerance of 0.25, apexTrackSNRThreshold of 1 and count thresholds of 1. Output was set to Apex3D csv file with -writeFuncCsvFiles to enable peak picking in all dimensions, including retention time (*t*_*R*_). For each run, the resulting csv file contains all peak picked ions with their apex in the *m/z*, drift time (*t*_*D*_) and *t*_*R*_ dimension, as well as their respective peak picking errors and the total summed intensity. In case of SONAR, *t*_*D*_ was mimicked by quadrupole selection within the Apex3D software and hence was processed identically. For all runs, subsequent analysis only retained HE fragment ions while LE precursor ions were discarded.

The single QC DDA run was converted to an mgf file with default parameters in Progenesis QI for Proteomics version 4.1.6675.48614 (Nonlinear Dynamics, Newcastle upon Tyne, UK).

#### 1.3 Experimental designs

Multiple experimental designs, i.e. experiments, were defined using different runs from different samples:

- 27 in-house HDMS^e^ runs: 9 runs per sample from condition A, B and QC
- 27 in-house SWIM-DIA runs: 9 runs per sample from condition A, B and QC
- 18 in-house HDMS^e^ and SWIM-DIA runs: 9 runs per acquisition, all from the same QC sample
- 1 in-house DDA run: 1 run from a QC sample
- 10 public HDMS^e^ runs from PXD001240: 5 runs per sample from condition A and B
- 9 public SONAR runs from PXD005869: 3 runs per amount of 0.5*µ*g, 1.0*µ*g and 1.5*µ*g, all from the same HeLa sample

Throughout the subsequent sections of this supplementary note, each experiment is analyzed independently from the others. Thus, any reference to *all runs* is assumed to mean all runs within a single experiment. For each single experiment, the csv files with peak picked ions from all its runs are simultaneously imported in a Python 3.6.6 environment to obtain a single dataset with all ions concurrently. Herein, each ion has the following descriptive attributes 1-3) the *m/z, t*_*D*_ and *t*_*R*_ apex and 4) run origin. Supporting attributes initially include 5-7) the error on the *m/z* (in parts per million (ppm)), *t*_*D*_ and *t*_*R*_ and 8) intensity. Throughout the creation of an ion-network, the additional supporting attributes 9-12) between-run calibrated *m/z, t*_*D*_, *t*_*R*_ and intensity and 13) aggregate index are appended.

#### 1.4 Between-run calibration

To calibrate the *m/z, t*_*R*_ and *t*_*D*_ of each run, the 50,000 most abundant ions of each run were selected. Since the *m/z* of all ions was already normalized post-acquisition by the lockmass throughout the Apex3D peak picking, this is generally the most accurate descriptive attribute of an ion. As such, the *m/z* distance (in ppm) was used as metric to perform a hierarchical agglomerative clustering with single linkage on all these ions. All clusters that contain each run exactly once were retained and considered potentially aligned prior to *t*_*R*_ and *t*_*D*_ outlier removal.

For each cluster the maximum distance in *t*_*R*_ and *t*_*D*_ between its constituent ions was calculated. Based on the distribution of the absolute deviation to the median of all *t*_*R*_ or *t*_*D*_ errors, individual *z*-scores were calculated per cluster. Each cluster with a *z*-score exceeding 5 was considered an outlier and removed. This process of outlier removal was repeated until only clusters with *z*-scores below 5 for both *t*_*R*_ and *t*_*D*_ remained. The final set of clusters was considered to be correctly aligned and equally partitioned into a set of clusters for calibration and validation. Note that the partitioning was done by selecting even and uneven clusters after *m/z* sorting, potentially introducing some dependency bias through isotopes between calibration and validation clusters.

For each calibration cluster, the average *m/z*, and *t*_*D*_ was calculated. Per run, the median error of its aligned ions towards their respective cluster average was calculated for the *m/z* (in ppm) and *t*_*D*_. These median run errors were subtracted from the original *m/z* (in ppm) and *t*_*D*_ to calibrate the *m/z* and *t*_*D*_ of all ions in the complete dataset.

To calibrate the *t*_*R*_ between runs, calibration clusters were first partitioned in multiple groups by a total order relation. More precise, for each pair (*a, b*) of calibration clusters from different groups and each run *s*, the *t*_*R*_ of the constituent ions always satisfies *t*_*R*_(*a*_*s*_) < *t*_*R*_(*b*_*s*_). Vice versa, for each calibration cluster *a* in a group containing multiple calibration clusters, there always exists a calibration cluster *b* in the same group and two runs *s* and *r* such that *t*_*R*_(*a*_*s*_) < *t*_*R*_(*b*_*s*_) while *t*_*R*_(*a*_*r*_) > *t*_*R*_(*b*_*r*_). Next, per group the average *t*_*R*_ of all constituent ions of all calibration clusters was determined per run as well as for all runs combined. Notice that both the group averages and group run averages always have the same ordering by definition of the total order relation. Finally, a calibration function was defined per run by applying a piece-wise linear transformation between run group averages and total group averages throughout the complete LC gradient (pseudo-groups defined for *t*_*r*_ = 0 and max(*t*_*r*_)). With this calibration function the *t*_*R*_ of all ions in the complete dataset were calibrated.

Finally, the validation clusters were used to obtain an automated estimate of the between-run errors of the calibrated *t*_*R*_ errors. Per validation cluster, the maximum distance of the calibrated *t*_*R*_ of its constituent ions was determined. The standard deviation of this distribution was considered the maximum error between two ions. Note that ions with larger errors can still be aligned, as long as there exists a path of pairwise alignment connecting them through intermediate runs.

#### 1.5 Ion-network generation

An ion-network was generated based on all calibrated ions in the dataset. First, ions are aligned into aggregates. The aggregates that contain reproducible ions comprise the nodes within this ion-network, while irreproducible ions are considered noise. Second, edges are set between aggregates where all constituent ions show consistent within-run co-elution. This deconvolutes HE fragment ions from chimeric precursors based on minor stochastic differences between runs, while HE fragment ions from the same precursor are connected with an edge (Supplementary Figure SF4).

##### 1.5.1 Nodes: between-run alignment and denoising

To align ions into aggregates, all ions in the entire experiment are considered concurrently. For each pair (*a, b*) of ions from two different runs, they are defined to be pairwise aligned if and only if their respective differences *d* in *m/z* (in ppm), *t*_*D*_ and *<u>t</u>*_*R*_ are within certain limits. For the *m/z* and *t*_*D*_, these limits satisfy 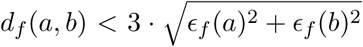 for {*f* ∈*m/z, t*_*D*_}with *E* their respective apex errors. For the *t*_*R*_, this maximum distance was determined previously by the validation clusters in the calibration step.

Once all ions are pairwise aligned, multiple consecutive pairwise alignments connect a set of ions into a cluster. While pairwise alignment is always between ions from different runs, consecutive pairwise alignments can connect multiple ions from the same run into a single cluster. For such a cluster, an aggregate cannot be defined as its constituent ions would be ambiguous. Therefore, a trimming is essential to remove all consecutive pairwise alignments connecting two ions from the same run. In a first step, each cluster that contains more than one ion per run is selected for trimming. For such a cluster, all non-transitive pairwise alignments are removed, i.e. for each retained pairwise alignment between ion *a* and *b* there exists an ion *c* such that both (*a, c*) and (*b, c*) are also pairwise aligned. If this step partitions a cluster into smaller clusters, each of these smaller clusters are again subjected to step one. Otherwise, the remaining pairwise alignments are trimmed by iteratively checking consecutive pairwise alignments of increasing length. Per iteration, it was checked whether there exists a consecutive pairwise alignment connecting two ions from the same run. If one or more of such consecutive pairwise alignments exists, all pairwise alignments in such consecutive connections are removed. If this iteration partitions a cluster in smaller clusters, each of these smaller clusters is again subjected to step one, otherwise the next iteration commences. By design, this process finishes at the latest after as many iterations as there are runs. Hereafter, no clusters containing multiple ions from the same run remain and all clusters can form aggregates with unambiguously aligned constituent ions.

As this trimming is quite stringent, a last step is performed which merges clusters not containing ions from the same run. This is done by iterating over all original untrimmed pair-wise alignments in order by Euclidean distance, i.e. a pairwise alignment defines a distance 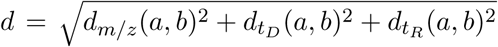. Once no clusters can be merged anymore, all clusters are defined as aggregates. Finally, all aggregates with reproducibility of at least two are defined as nodes in the ion-network.

##### 1.5.2 Edges: consistent within-run co-elution and deconvolution

An edge is set between two aggregates *a* and *b* if and only if they consistently co-elute. Two aggregates are defined as consistently co-eluting if and only if their constituent ions co-elute in each overlapping run, i.e. for each run *s* with ions *a*_*s*_ and *b*_*s*_ we have 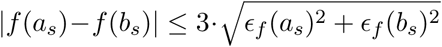 for *f ∈* {*t*_*R*_, *t*_*D*_} with *f*_*ϵ*_ the estimated apex error.

However, a large run count can introduce a dimensionality curse, meaning that the constituent ions of two aggregates from the same precursor can have peak picking errors that by chance are too large to define co-elution in some run. Therefore the definition of consistently co-eluting is weakened to mean that they should co-elute in at least 90% of the runs where they both are found (rounded down). As a final constraint, two aggregates need to co-elute in at least two runs to be considered *consistently* co-eluting.

#### 1.6 Intensity normalization

To normalize intensity differences between all runs from all samples, the average intensity of all fully reproducible aggregates was calculated. Next, the logFC distance to the average of each constituent ion of these fully reproducible aggregates was determined per run. For each run, the median of these logFC distances were determined and subsequently subtracted (in logarithmic space) from all ions in the complete dataset.

#### 1.7 Database search

Once a proteomic ion-network has been created, it can be annotated. A fasta file containing all SwissProt entries from Human, Yeast and E. coli was downloaded (November 23, 2018) for all but the SONAR experiment that only used Human. The common repository of adventitious proteins (cRAP) database was appended, as well as decoys containing all reversed protein sequences. A standard in silico tryptic digest without missed cleavages was performed. Fixed modification of cysteine was set to +57.021464 (carbamidomethyl) and no variable modifications were considered. Duplicate peptides from different proteins were merged to obtain a list of unique peptide sequences. Peptides originating solely from decoy proteins were classified as decoy peptides, while all others were classified as targets. All fragments, i.e. mono-isotopic masses of all singly-charged b- and y-ions, were calculated for each peptide.

For each aggregate that has at least two other consistently co-eluting aggregates, all potential fragment explanations *p*_1_, …, *p*_*n*_ were determined within 20 ppm of its *m/z*. For each of these fragment explanations *p*_*i*_, the count *c*_*i*_ of consistently co-eluting aggregates with a fragment explanation covering the same peptide was determined. Hereafter, the logarithmic cumulative frequency *f*_1_, …, *f*_*m*_ of fragment explanations *p*_*i*_ with count *c*_*i*_ ≥1, …, *m* was determined. A robust linear regression *r* was performed by random sample consensus (RANSAC) for all but the latest logarithmic cumulative frequencies *f*_1_, …, *f*_*m−*1_. Roughly interpreted, a score *r*(*i*) coincides with an *e*-value describing the likelihood that this count *c*_*i*_ is a random event. Finally, this linear regression was extrapolated to the point *m* and all fragment explanations *p*_*i*_ with count *c*_*i*_ = *c*_*m*_ were given the score *s* = −*r*(*m*). Each of these particular fragment explanations *p*_*i*_ of an aggregate are hereafter defined as a peptide-fragment-to-ion-neighborhood match (PIM), analogous to a precursor that is assigned a peptide-to-spectrum match (PSM) in DDA. Note that not all aggregates are given a PIM, as there sometimes are no fragment explanations or no linear regression can be made due to too few consistently co-eluting aggregates. Equally, some aggregates are assigned more than one PIM, which by the current definition always have an equal score.

As an additional accuracy measure besides a PIM score, the *t*_*D*_ of each aggregate was used as a proxy for potential precursor *m/z*. First, the aggregates of each PIM were checked for a consistently co-eluting aggregate with an unfragmented singly, doubly or triply charged precursor *m/z* of the covering peptide within 20 ppm. For each of these selected PIMs and per charge state, a linear regression is then made by RANSAC so that the theoretical *m/z* can be predicted in function of *t*_*D*_ for all three charges. Finally, three differently charged precursor *m/z* values are predicted for each aggregate based on its *t*_*D*_ and the distances between these three theoretical *m/z* values and the potential *m/z* values of the PIMs precursor are determined. The minimum of these three distances (in standard deviations) is taken to set the most likely precursor charge of a PIM.

Each PIM is then rescored and assigned a target-decoy false discovery rate (FDR) controlled *q*-value by percolator [4], in which they are treated as traditional PSMs. The flags “-D 15” to use all documented features, “-I concatenated” as decoy and “-A” to use fido algorithm for protein scoring [5], as well as the following features per aggregate are passed to percolator:

- *t*_*R*_
- Fragment explanation *m/z* difference (in ppm)
- Aggregate reproducibility
- Number *k* of consistently co-eluting aggregates
- Count *c*_*m*_ of consistently co-eluting aggregates with fragment explanations covering the same peptide
- Consistently co-eluting aggregate match ratio 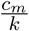
- Estimated precursor charge *z*
- Number of standard deviations between peptide *m/z* and predicted precursor *m/z*
- Peptide length *l*
- Peptide match ratio 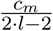
- Score *s*

#### 1.8 Interactive graphical browser

After creating and annotating an ion-network, it can be visualized in an interactive graphical browser. Interactive options include:

- Function to zoom into the region of interest (*t*_*R*_ and *t*_*D*_ coordinates)
- Show only those aggregates that satisfy a selected reproducibility
- Turn on/off edges between consistently co-eluting aggregates
- Label aggregates by their peptide annotation, protein annotation, calibrated *m/z*, calibrated *t*_*R*_ or calibrated *t*_*D*_
- Set FDR threshold to color and label annotated aggregates
- Select aggregates to show the logarithmic calibrated intensity, uncalibrated *t*_*R*_ or uncalibrated *t*_*D*_ of its constituent ions
- Export currently visible aggregates or ions to a variety of image formats (png, svg, pdf, ….)

### 2 Availability and reproducibility

In accordance with the European Bioinformatics Community (EuBIC) guidelines (https://eubic.github.io/ReproducibleMSGuidelines/), all data and software are made publicly available to ensure full transparency and reproducibility. The only exception is the peak picking software Apex3D (Waters Corporation), that is not freely available. A freely available alternative to Apex3D peak picking is IMTBX [6], but this has not yet been tested for compatibility.

#### 2.1 Data

All data are available at ProteomeXchange (PXD015318). This includes raw data and acquisition parameters for in-house runs as well as all peak picked data, since Apex3D is not freely distributed.

All files to (re)create and (re)analyze ion-networks for all experiments (including parameters, logs, figures and (intermediate) results) are deposited alongside this data.

QC files monitoring general MS performance are also included under the same ProteomeXchange identifier.

#### 2.2 Software

The complete source code (version 0.1.190809) to create and analyze ion-networks is available at GitHub (https://github.com/swillems/histopya), including download/installation instructions and custom scripts only used within this manuscript. Full reproduction of all results, including figures, is possible but requires external files to be downloaded from ProteomeXchange due to size limitations. This GitHub repository includes a minor tutorial test case to illustrate how to use the software on a novel experiment provided by the user.

All analyses in this manuscript were performed on a CentOS Linux release 7.6.1810 (Core) with 88 (after hyperthreading) CPUs (Intel(R) Xeon(R) Gold 6152 CPU @ 2.10GHz) and 754 Gb RAM, but less powerful systems suffice for all experiments that are presented here. Peak picking was done through wine64 version 4.0 (with prior taskset -c option to use only 44 CPUs to circumvent the maximum 64 CPU restriction) as Apex3D is a windows executable.

## Acronyms

*t*_*D*_: drift time
*t*_*R*_: retention time
*m/z*: mass-to-charge ratio
cRAP: common repository of adventitious proteins
CV: coefficient of variation
DDA: data-dependent acquisition
DIA: data-independent acquisition
diaPASEF: parallel accumulation – serial fragmentation combined with data-independent acquisition
FDR: false discovery rate
HDMS^**e**^: high definition Ms^e^
HE: high energy
IMS: ion mobility separation
IQR: interquartile range
LC: liquid chromatography
LE: low energy
logFC: logarithmic fold change
MRM: multiple-reaction-monitoring
MS: mass spectrometry
PIM: peptide-fragment-to-ion-neighborhood match
ppm: parts per million
PSM: peptide-to-spectrum match
QC: quality control
RANSAC: random sample consensus
SNR: signal-to-noise ratio
SONAR: scanning quadrupole DIA
SWATH: sequential window acquisition of all theoretical mass spectra
SWIM: single window ion mobility
ToF: time of flight
XIC: extracted ion chromatogram

